# Biological Control of Green Mold in Simulated Post-harvest Chain of Citrus Fruit: Efficacy of *Candida oleophila* Strain O and Molecular Insight into Elicitation of Host Immune System

**DOI:** 10.1101/2024.03.05.583495

**Authors:** Ermes Ivan Rovetto, Federico LA Spada, Soumia EL Boumlasy, Sebastiano Conti Taguali, Mario Riolo, Antonella Pane, Santa Olga Cacciola

**Affiliations:** Department of Agriculture, Food and Environment, University of Catania, 95123 Catania, Italy; Laboratory of Materials-Catalysis, Chemistry Department, Faculty of Science, University Abdelmalek Essaadi, Tetouan B.P. 2117, Morocco; Department of Agricultural Science, Mediterranean University of Reggio Calabria, 89122 Reggio Calabria, Italy

**Keywords:** *Candida oleophila*, citrus green mold, *Penicillium digitatum*, citrus fruit supply chain, biological control, plant defense transcriptomic response

## Abstract

Managing post-harvest decays in citrus fruit without relying on conventional pesticides presents a significant challenge in modern Plant Pathology. This study aimed to evaluate the efficacy of the biological control agent *Candida oleophila* strain O in controlling green mold caused by *Penicillium digitatum* throughout various stages of the post-harvest supply chain. Using a series of *in vivo* experiments, different scenarios of *P. digitatum* infections in clementine tangerine, orange, and lemon fruit were examined, with treatments applied before, during or after infection. The study simulated typical conditions of the citrus supply chain, including picking, processing in packinghouses, and transportation, as well as cold storage and shelf-life phases. Results indicated that *C. oleophila* exhibited significant efficacy in reducing green mold symptoms, even at shelf-life temperatures, making it a practical alternative to conventional fungicides. Additionally, the study provided insights into the molecular mechanisms underlying the defensive response of citrus fruit to *C. oleophila* treatment, with up-regulation of defense-related genes observed across different fruit types. Overall, this study underscores the potential of *C. oleophila* as a sustainable and effective solution for managing post-harvest decays in citrus fruit within the complexities of the supply chain.

## 1. Introduction

The primary and globally prevalent postharvest challenge impacting citrus fruit is the green mold caused by *Penicillium digitatum* (Abo-Elnaga, 2013; Kanan and Al-Najar, 2008; Plaza et al., 2004; Rovetto et al., 2023, 2024), a destructive disease accounting approximately 90% of total post-harvest losses (Costa et al. 2019; Ismail and Zhang, 2004). This disease causes considerable damages on major commercial citrus crops, specifically oranges (Elsherbiny et al., 2021), lemons (Ben-Yehoshua et al., 2008), tangerines (Suwannarach et al., 2015), and grapefruit (Shi et al., 2018). The citrus green mold is caused by a wound-mediated infection of the fruit peel (Rovetto et al., 2024). At early stages, the infection appears as a softened area of the peel surrounding the infection point (La Spada et al. 2021); this symptom, known as ’Clear Rot’ (Ismail and Zhang, 2004), creates conducive conditions for the development of white mycelium and triggers the rapid decay of fruit. Within a few days, the infection progresses to complete molding of the fruit. Simultaneously, the pathogen develops a substantial quantity of green airborne conidia on the fruit surface, serving as a source of inoculum for new infections (Costa et al., 2019; Ismail and Zhang, 2004; Rovetto et al., 2024; Vu et al., 2018).

Due to the high virulence of *P. digitatum* and the potential for related infections to occur at any point, from the field to the consumers’ table, the post-harvest supply chain of citrus fruit—encompassing phases of harvesting, warehouse processing (including treatments and packaging), transportation, and delivery to the final customer—deserves particular attention and the implementation of specific practices of control (Fallanaj et al., 2016). Therefore, care in both harvesting and handlings fruit as well as sanitization measures in packing-houses are effective in preventing the disease to minimize peel punctures, wounds, bruises, compression damages, and other mechanical (El-Otmani et al., 2011; Zacarias et al., 2020). Respecting specific temperatures after the warehouse processing, such as 4-8°C during shipment, is an additional practice for limiting pathogen’s development in the short term (Khumalo et al., 2021). While these practices could be easily applicable in short-duration supply chains, such as local or national fruit markets, their applicability faces technical limitations in continental and transcontinental markets, where long-duration shipments of fruit at a continuous cold temperature are not cost effective and, therefore, cannot be guaranteed. In these latter contexts, the major strategy for limiting losses by citrus green mold relies prevalently on post-harvest treatments of fruit with synthetic fungicides (Ismail and Zhang, 2004). In this respect, the widely employed fungicide imazalil (IMZ), well known for its strong effectiveness against *Penicillium* spp., provides both advantages and drawbacks (Bus et al., 1991, 1992; Davé et al., 1989; Holmes and Eckert, 1999; Ismail and Zhang, 2004; Moraes Bazioli et al., 2019; Sánchez-Torres and Tuset, 2011; Smilanick et al., 2005). Indeed, while IMZ application is essential for maintaining fruit quality, its employment has several drawbacks, including the environmental impact, the presence of toxic residues on the fruit, that in the EU countries is regulated by stringent legislative measures, and the risk of developing resistant pathogen isolates, consequently diminishing its efficacy (Arslan, 2015; Davé et al., 1989; EFSA, 2010; Gikas et al., 2022; Holmes and Eckert, 1999; Kanetis et al., 2007; Stracquadanio et al., 2021; Strange and Scott, 2005). Consequently, this has fostered the development of eco-friendly disease management strategies for controlling citrus green mold, that can be an alternative to synthetic fungicides or may contribute to reduce their use (Fenta et al., 2019; Hammami et al., 2022; Janisiewicz and Korsten, 2022; La Spada et al., 2021; Li et al., 2022; Riolo et al., 2023; Stracquadanio et al., 2020; Sui et al., 2016; Talibi et al., 2014). These include, among others, synthetic resistance elicitors, that activate natural plant defense mechanisms, and biological control agents (BCAs) (Battha, 2022; Boller, 1995; Côté et al., 1998; Cluzet et al., 2004; Droby et al., 2009; Du Jardin, 2015). Among resistance elicitors, salts of phosphonic acid (phosphonates) have gained attention for their applications in agriculture as either fertilizers or fungicides (Gómez-Merino and Trejo-Téllez, 2015; Manghi et al., 2021). Presently, phosphonates are banned from organic agriculture by the EU and are restrictively classified as plant protection products (European Parliament, 2019). Although phosphonates have been exploited mainly for controlling plant diseases caused by Oomycetes, potassium phosphonate was reported to be also effective against green mold of citrus fruit (Amiri and Bompeix, 2011; Cerioni et al., 2013; Strano et al., 2015; Souza Morais Reis et al., 2024). Among potential BCAs of post-harvest rots of citrus fruit, extensive research has been conducted on various species of yeasts, including *Candida* spp, *Cryptococcus* spp., *Metschnikowia* spp., *Pichia* spp. and *Rhodotorula* spp. (Droby et al., 1993; Hammami et al., 2022; Kowalska et al., 2022; Liu et al., 2013; Oztekin et al., 2023; Zhang et al., 2020). *Candida* species, particularly *Candida oleophila*, are at the forefront of commercial biopesticides, due to their positive effects on plants both as biological control agent and elicitor of plant defense response (Kowalska et. al., 2022). Currently, the strategic application of elicitors of plant defense response is emerging as a promising avenue for fortifying disease resistance in agronomic crops (Boller, 1995; Cluzet et al., 2004; Côté et al., 1998; La Spada et al., 2021). In this respect, it is well known that plants respond to infections through distinct biochemical changes, which are mediated by the transcription of various defense genes both locally and distantly from the infection site (Boava et al., 2011). These responses encompass the hypersensitive response (HR), modifications in the cell wall, production of reactive oxygen species (ROS), accumulation of phytoalexins, and synthesis of pathogenesis-related (PR) proteins (Boava et al., 2011; La Spada et al., 2021; Van Loon et al., 2006). Some PR proteins, such as β-1,3-glucanases, can degrade fungal cell wall constituents, specifically β-1,3-glucan and chitin, exhibiting antifungal properties (Balasubramanian et. al., 2012); for this reason, their biosynthesis and accumulation are considered a fundamental defense mechanism (Odjakova and Hadjiivanova, 2001). Other PR proteins, such as peroxidases play, an additional key role in the plant defense response; produced during lignin synthesis, they contribute to reinforcing the cell wall, thereby enhancing resistance against numerous pathogens (Miedes et al., 2014). The antioxidant activity of peroxidase has been documented for its positive correlation with the enhancement of resistance of citrus fruit to *Penicillium* (Ballester et al., 2006). In addition to PR-proteins, the phenylalanine ammonia-lyase (PAL), the first enzyme in the phenylpropanoid pathway, increases the plant defense response to various types of stress (Ke and Saltveit, 1989; Youssef et al., 2014; Wang et al., 2023). In citrus fruit, the activity of phenylalanine ammonia-lyase (PAL) leads to the production of flavonoids (Arcas et al., 2000; Del Río et al., 1998) and p-coumaric acid derivatives (Afek et al., 1999; Kim et al., 1991), potentially bolstering the fruit’s protection against pathogens. Several studies have delved into evaluating the antagonistic activity of *C. oleophila* for controlling citrus green mold (Dukare et al., 2019; Laahlali et al., 2004; Lassois et al., 2008). Additionally, a study of the 2002 by Droby et al. evaluated the ability of *C. oleophila* in affecting the plant biosynthesis of some defensive metabolites in peels of grapefruit under *P. digitatum* infection. Despite this attention, there remains a notable gap in understanding the effectiveness of *C. oleophila* in controlling citrus green mold throughout the various stages of the citrus fruit supply chain. Moreover, there is limited knowledge concerning *C. oleophila*’s ability to modulate the triggering of specific genetic pathways related to the plant-defense response, and whether this ability varies depending on the type of citrus fruit.

To advance beyond the current state of the art, this study simulated different scenarios of *P. digitatum* infections in various types of citrus fruit (clementine tangerines, oranges, and lemons) to assess the potential of *Candida oleophila* in reducing the incidence of citrus green mold throughout the different stages of the citrus fruit supply chain. Additionally, the study evaluated *C. oleophila*’s ability to modulate the plant-defense response by analyzing the transcription of genes encoding β-1,3-glucanases, phenylalanine ammonia-lyase, and peroxidase in clementine tangerine, orange, and lemon fruit.

## 2. Materials and Methods

### 2.1 Fungal plant pathogen

The IMZ-sensitive (MIC < 1 μml^−1^ a.i.) *P. digitatum* strain P1PP0 used in this study was from the collection of the Laboratory of Molecular Plant Pathology of the Department of Agriculture, Food and Environment of the University of Catania (Catania, Italy) and was characterized in a previous study (El boumlasy et al., 2021). Before tests, the *P. digitatum* strain P1PP0 was maintained on Potato Dextrose Agar at 25°C.

### 2.2. Biocide agent and experimental substances

The substances used in this study were: a 10^8^ CFU·mL^-1^ suspension in sterile distilled water (sdw) of the yeast *Candida oleophila* Montrocher strain O (ID substance I01), belonging to the collection of the Molecular Plant Pathology Laboratory of the Department of Agriculture, Food and Environment (University of Catania, Catania, Italy); an eco-friendly sanitizing agent containing hydrogen peroxide and acetic acid (ID substance P01); a 57% (w/v) potassium phosphonate formulation (ID substance RI01); and an emulsifiable 50% w/v concentrate formulation of imazalil fungicide (ID substance F01). All chemicals used in this study were kindly provided by DECCO Italia srl (Belpasso, CT, Italy), and their application dosages are reported in Table 1.

**Table 1.**
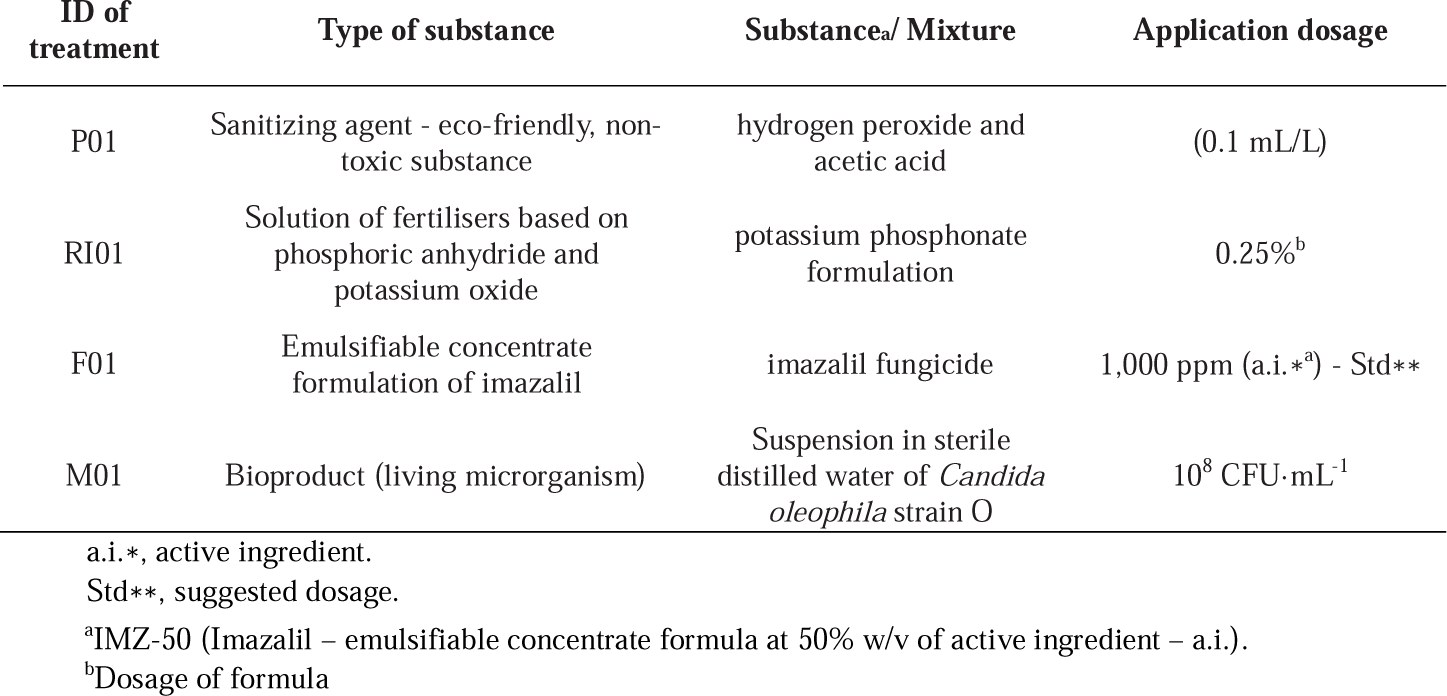
List of substances used in this study.

### 2.3. Citrus fruit

Fruit used in this study were mature and non-wounded clementine tangerines (*Citrus clementina* Hort. ex Tan., cv. Comune), oranges (*Citrus x sinensis* (L.) Osbeck, cv. Ovale) and lemons (*Citrus x limon* (L.) Burm. f., cv. Femminello) collected from an organically cultivated orchard. Before each experiment, fruit underwent preliminary disinfection through a sequence of processes, including washing in NaClO 1%, followed by rinsing in sterilized distilled water (sdw), and subsequent air drying at room temperature.

### 2.4. Evaluating the effectiveness of *Candida oleophila* strain O in controlling citrus green mold

The effectiveness of *Candida oleophila* strain O (M01) in controlling citrus green mold was assessed in orange, lemon, and clementine tangerine fruit. Tests were conducted to simulate as close as possible various stages of the citrus postharvest chain, spanning from warehousing (including treatment, packaging, and storage) to retailing. For each type of fruit, these stages were replicated through two consecutive phases following treatment: a. 7-day storage under refrigerated conditions (at 6°C) and b. an additional 7-day storage under shelf-life conditions (at 18°C). These storage conditions were included within three realistic scenarios of fruit infection by *P. digitatum*: i. before treatment; ii. during treatment; iii. 12 hours post-treatment. Further details of the conducted tests are provided in subsequent sections.

#### 2.4.1. Fruit inoculated with *P. digitatum* before treatment

The peel of each citrus fruit was wounded by a plastic tip (diameter, 2 mm) at four equidistant points in an equatorial position along the rind surface. Then, each wound was inoculated with 5 μl of a conidial suspension (10^6^ conidia/ml) of *P. digitatum* strain P1PP0; fruit were then incubated at 20°C for 4 hours. After the incubation period, inoculated fruit were pre-treated by dipping in P01 (0.1 mL/L) for 30 seconds and rinsing in sdw for additional 30 seconds, or by a simple dipping in sdw for 30 seconds (control). Immediately after the pre-treatment, at each wound, fruit were treated with a 65 μl spray of RI01, F01, M01 or sdw (controls). Then, fruit were organized in five batches corresponding to the following treatments: i. fruit pre-treated only with sdw (control 1), ii. fruit just pre-treated only with P01 (control 2), iii. fruit pre-treated with P01 and then treated with RI01, iv. fruit pre-treated with P01 and then treated with F01, v. fruit pre-treated with P01 and then treated with M01. Each treatment comprised three technical replicates made up of five citrus fruit each.

Fruit from each treatment were air dried, packaged in plastic bags, then incubated at 6°C for 7 days. After the incubation period, fruit were further incubated at 18°C for other 7 days. For each treatment, the disease incidence, evaluated as occurrence of disease symptoms (%), was recorded at 7, 10 and 14 days post the last treatment session.

The above described experimental scheme was replicated for orange, lemon and clementine tangerine fruit.

#### 2.4.2. Fruit inoculated with *P. digitatum* during treatment

The peel of each citrus fruit was wounded by a plastic tip (diameter, 2 mm) at four points along the equatorial surface. Then, fruit were pre-treated by dipping in P01 (0.1 mL/L) for 30 seconds and rinsing in sdw for additional 30 seconds, or by a simple dipping in sdw for 30 seconds (control). After the pre-treatment, each wound was sprayed with 65 μl of RI01, F01, M01 or sdw (controls); then, each wound was inoculated with 5 μl of a conidial suspension (10^6^ conidia/ml) of *P. digitatum* strain P1PP0. Fruit were air dried and organized in five batches corresponding to the following treatments: i. fruit pre-treated only with sdw (control 1), ii. fruit pre-treated only with P01 (control 2), iii. fruit pre-treated with P01 and then treated with RI01, iv. fruit pre-treated with P01 and then treated with F01, v. fruit pre-treated with P01 and then treated with M01. Each treatment comprised three technical replicates each with five citrus fruit. After drying, fruit from each treatment were packaged in plastic bags and incubated at 6°C for 7 days. After the incubation period, fruit from any treatment were further incubated at 18°C for other 7 days. For each treatment, the disease incidence, evaluated as occurrence of disease symptoms (%), was recorded at 7, 10 and 14 days post the last treatment session.

The above-described experimental scheme was replicated for orange, lemon and clementine tangerine fruit.

#### 2.4.3. Fruit inoculated with *P. digitatum* post-treatment

The peel of each citrus fruit was wounded by a plastic tip (diameter, 2 mm) at four points along the equatorial surface. Then, fruit were pre-treated by dipping in P01 (0.1 mL/L) for 30 seconds and rinsing in sdw for additional 30 seconds, or by a simple dipping in sdw for 30 seconds (control). After the pre-treatment, each wound was sprayed with 65 μl of RI01, F01, M01 or sdw (controls); then, each wound of each fruit was inoculated with 5 μl of a conidial suspension (10^6^ conidia/ml) of *P. digitatum* strain P1PP0. Fruit were air dried and organized in five batches corresponding to the following treatments: i. fruit pre-treated only with sdw (control 1), ii. fruit pre-treated only with P01 (control 2), iii. fruit pre-treated with P01 and then treated with RI01, iv. fruit pre-treated with P01 and then treated with F01, v. fruit pre-treated with P01 and then treated with M01. Each treatment had three technical replicates made up of five citrus fruit each. After drying, fruit from each treatment were packaged in plastic bags and incubated at 6°C for 12 hours. After the incubation period, each wound of any fruit was inoculated with 5 μl of a conidial suspension (10^6^ conidia/ml) of *P. digitatum* strain P1PP0. Fruit from all the treatments were packaged again and incubated at 6°C for additional 7 days. After the incubation period, fruit were further incubated at 18°C for other 7 days. For each treatment, the disease incidence, evaluated as occurrence of disease symptoms (%), was recorded at 7, 10 and 14 days post the last treatment session.

The above-described experimental scheme was replicated for orange, lemon and clementine tangerine fruit.

### 2.5. Tissue sampling for RNA isolation from fruit

For the evaluation of the differential expression of genes involved in the induction of resistance in citrus fruit (clementine tangerines, oranges and lemons) by the different treatments, a specific test was performed. Disinfected fruit were individually wounded with a 2 mm plastic tip at eight points on the rind surface in an equatorial position. The experimental design comprised six treatments as it follows: i. unwounded fruit; fruit treated with: ii. sdw; iii. P01; iv. RI01; v. F01; or vi. M01. For each treatment, fruit were collected at three different time points: 0, 24, and 48 h. For each time point, fruit batches comprised three fruit. Fruit of each treatment were placed in plastic containers and incubated at 18°C for 48 h. At each time point, fragments of peel (5 mm ×15 mm × 2 mm) were excised from each fruit of any treatment from the treated area. The excised tissues were rapidly frozen in liquid nitrogen and stored at −80°C until they were used for gene expression analyses. The experiment was repeated twice with similar results, so only results of the first experiment are reported.

#### 2.5.1 RNA extraction from fruit peels and synthesis of complementary DNA

Total RNA was extracted from 100 mg of frozen citrus fruit peels (clementine tangerines, oranges and lemons), which were reduced into a fine powder using liquid nitrogen. The RNeasy Plant Mini Kit (Qiagen, Venlo, Netherlands) was used for RNA extraction following the manufacturer’s protocol. Subsequently, the extracted RNA underwent treatment with the TURBO DNA-free™ Kit (Invitrogen, Carlsbad, CA, United States). The concentration of RNA was adjusted to 200 ng/μl, and its integrity was assessed through denaturing RNA electrophoresis in TAE agarose, as described by Masek et al. (2005). Reverse transcription was carried out using the High-Capacity cDNA Reverse Transcription Kit (Applied Biosystems™, Foster City, CA, United States) following the manufacturer’s instructions.

#### 2.5.2. Selection of genes and specific primers

A set of distinct primer pairs targeting various genes associated with induced resistance in citrus fruit were selected from previous studies (La Spada et. al., 2021; Youssef et al., 2014). Details about selected genes and related primers are reported in Table 2.

**Table 2.**
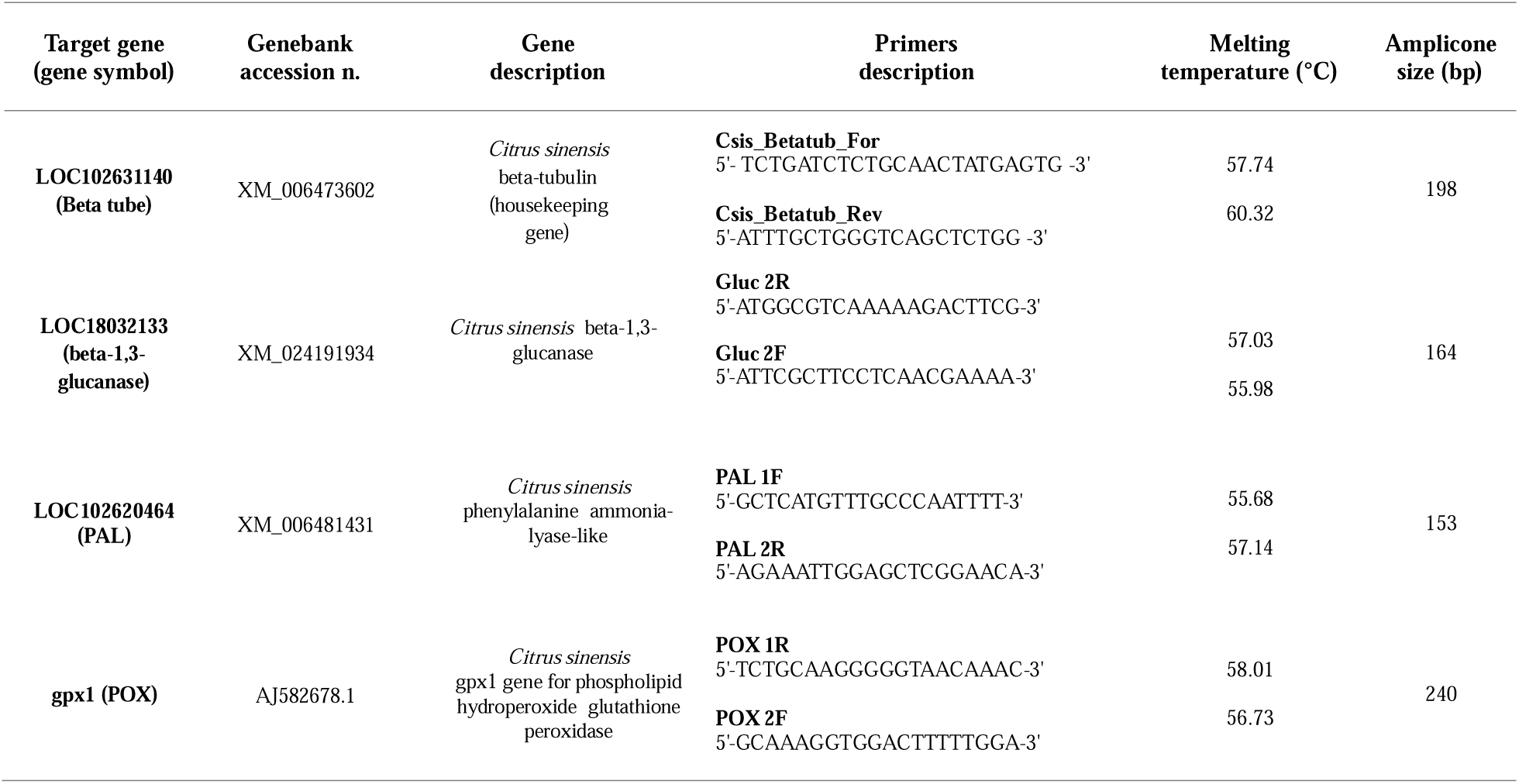
Gene-specific primers used in quantitative reverse transcription real-time polymerase chain reaction (qRT-PCR).

#### 2.5.3. Quantitative Real-Time PCR (qRT-PCR) analysis of gene expression

qRT-PCR amplifications were carried out by using the iCycler iQ™ Real-Time PCR Detection System (QuantGeene 9600, Hangzhou Bioer Technology Co., Ltd., BIOER, 1192 Bin An Rd, Binjiang District, Hangzhou, 310053, People’s Republic of China). Each reaction was carried out in a final volume of 20 μl, comprising 10 ng of cDNA, 1 μl of 10 μM for each primer, and 10 μl of PowerUp™ SYBR™ Green Master Mix (2X, Applied Biosystems™, Foster City, CA, United States). For each biological sample collected, qRT-PCR experiments were performed in triplicate. Thermocycling conditions were 2 min at 50°C (UDG activation), 2 min at 95°C (Dual-Lock™ DNA polymerase), followed by 40 cycles of two steps: 95°C for 15 s (denaturation) and 59°C (annealing/extension) for 1 min. The quantification of gene expression relative to the unwounded control sample was carried out by using the 2^-ΔΔCt^ method (Livak and Schmittgen, 2001), where ΔΔCt = (Ct of target gene − Ct of reference gene)_sample_ − (Ct of target gene − Ct of reference gene)_calibrator_ and Ct is the threshold cycle of each transcript, defined as the point at which the amount of amplified target reaches a fixed threshold above the background fluorescence.

### 2.5. Statistical analyses

Analysis of data was performed by one-way ANOVA followed by Tukey’s HSD (Honestly Significant Difference) post hoc test by using the R software. To normalize the distributions, percentage data were transformed into square root values, but untransformed percentages are reported in the respective graphs. Differences at p ≤ 0.05 were considered significant. The mean ± SD was reported in graphs.

## 3. Results

### 3.1. Effectiveness of *Candida oleophila* strain O in controlling citrus green mold

Following sections describe results on *in vivo* tests carried out to evaluate the effectiveness of the *C. oleophila* strain O in controlling citrus green mold on artificially *P. digitatum*-infected orange, lemon, and clementine tangerine fruit. In detail, three realistic scenarios of fruit infection were simulated: *P. digitatum* inoculation i. before treatment, ii. during treatment and iii. 12 hours post-treatment. Additionally, in order to simulate various stages of the citrus postharvest chain, per each scenario of inoculation the following stages of storing conditions were assessed: a. 7-day storage under refrigerated conditions (at 6°C), and b. an additional 7-day storage under shelf-life conditions (at 18°C).

#### 3.1.1. Effectiveness of *Candida oleophila* strain O on clementine tangerines

Results of tests on clementine tangerines are summarized in Fig.1. Seven days after the last treatment session, none of the clementine tangerine fruit from any treatment or infection scenario showed any symptoms of disease (Figure 1 - a1, b1, and c1). Ten days after the last treatment session, in any scenario of infection, clementine tangerines from treatment with M01 and F01 showed the lowest occurrences of disease symptoms (%), with statistical differences with water treated control; however, in any scenario of infection, the treatment with M01 determined an occurrence of disease symptoms (%) comparable to that of fruit from treatment with both RI01 and F01 (Figure 1 – a2, b2, and c2). A similar trend of occurrence of disease symptoms (%) was also observed fourteen days post the last treatment (Figure 1 – a3, b3, and c3).

**Figure 1.**
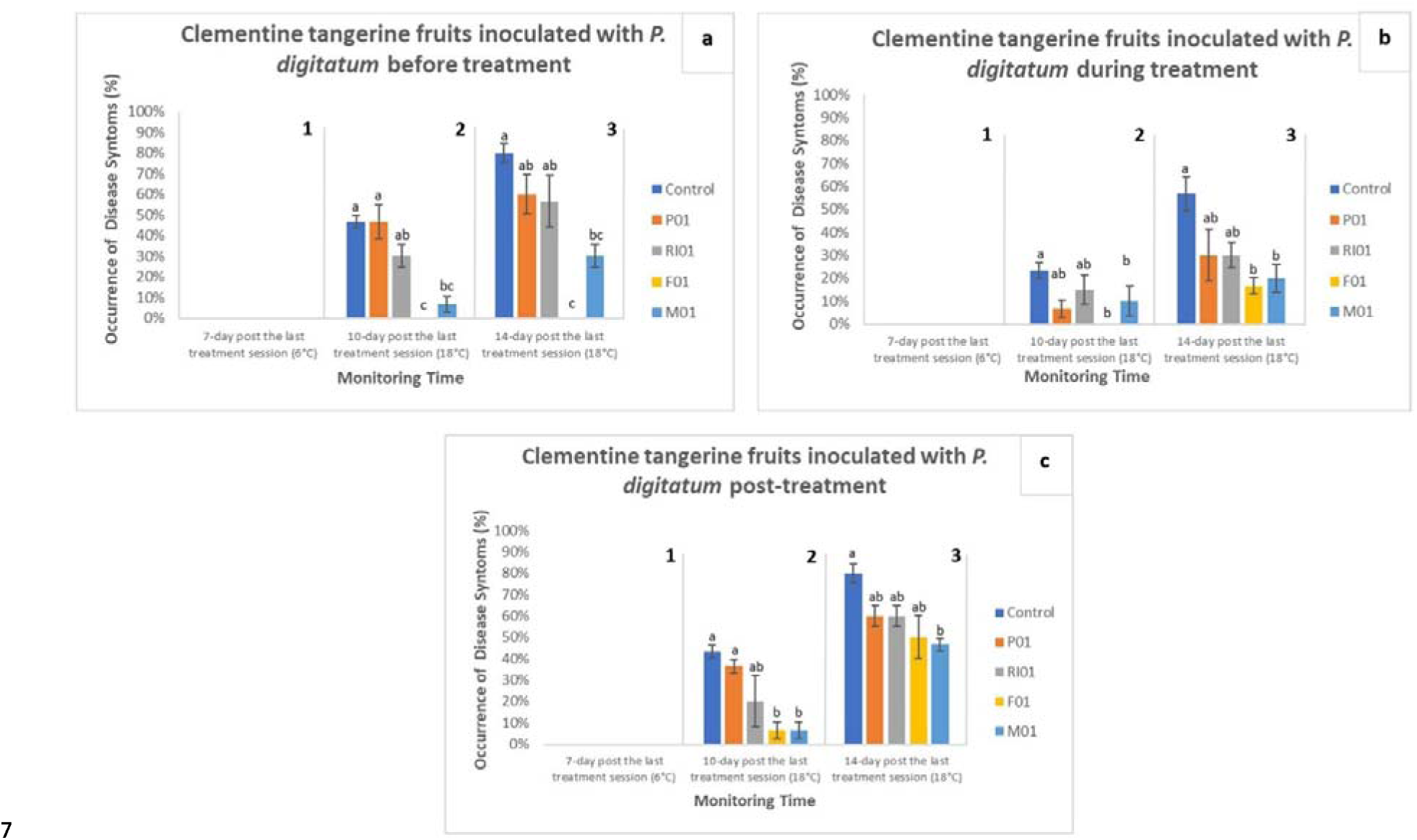
Evaluation of occurrence of disease symptoms (%) in clementine tangerine [*Citrus clementina* Hort. ex Tan.,cv. Comune] fruit in three scenarios of infection by *Penicillium digitatum* strain P1PP0: (a) before treatment; (b) during treatment and (c) 12 hours post-treatment. For each treatment, the occurrence of disease symptoms (%) was recorded at 7 (1), 10 (2) and 14 (3) days post the last treatment session. For each time point of measurement, values sharing the same letters are not statistically different according to the Tukey’s HSD (Honestly Significant Difference) test (p ≤ 0.05).

#### 3.1.2. Effectiveness of *Candida oleophila* strain O on oranges

Results of tests on oranges are summarized in Fig.2. As observed for clementine tangerines, seven days after the last treatment no orange fruit from any treatment or infection scenario showed any symptoms of disease (Figure 2 - a1, b1, and c1).

**Figure 2.**
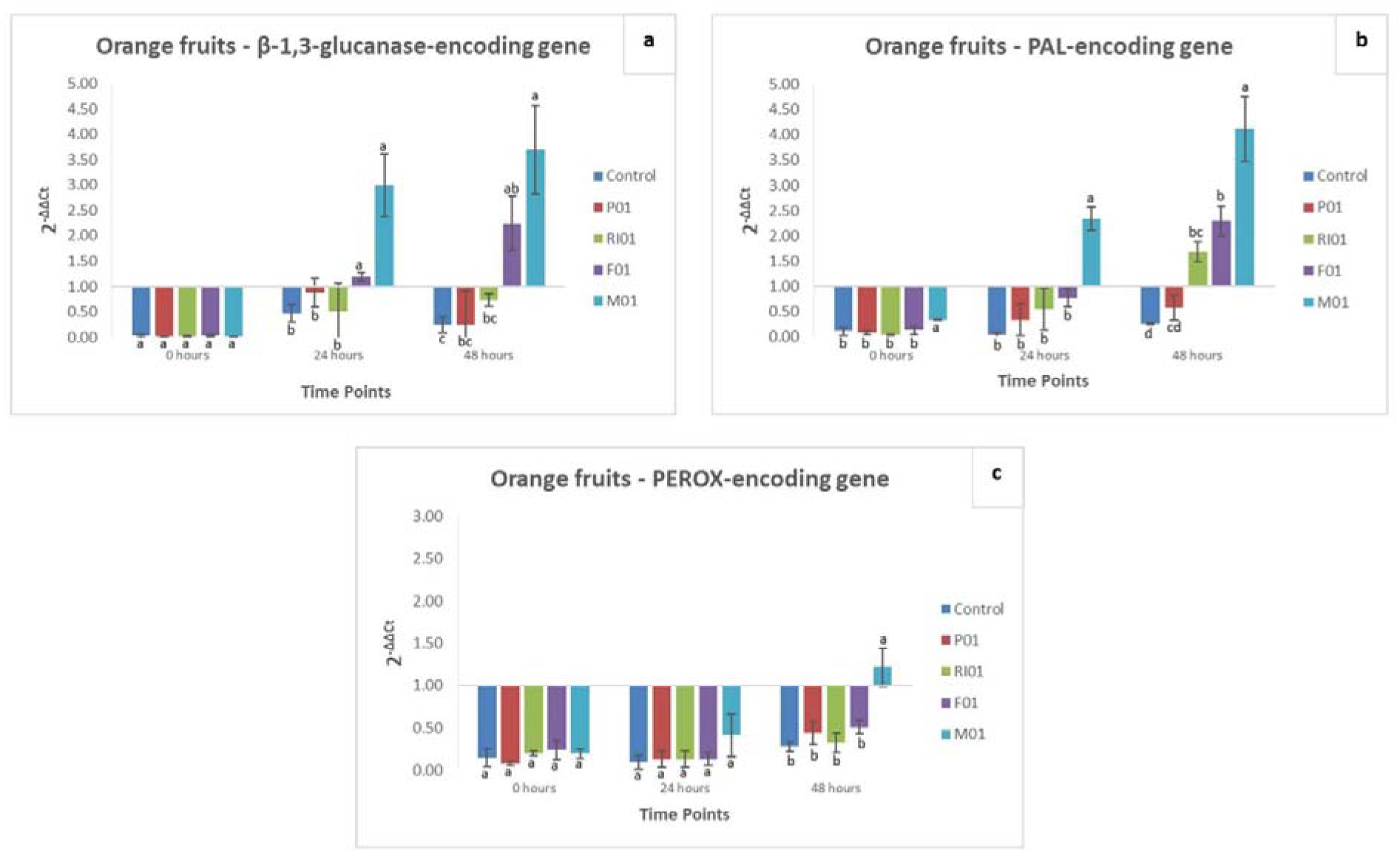
Evaluation of occurrence of disease symptoms (%) in orange [*Citrus x sinensis* (L.) Osbeck cv. Ovale] fruit in three scenarios of infection by *Penicillium digitatum* strain P1PP0: (a) before treatment; (b) during treatment and (c) 12 hours post-treatment. For each treatment, the occurrence of disease symptoms (%) was recorded at 7 (1), 10 (2) and 14 (3) days post the last treatment session. For each time point of measurement, values sharing the same letters are not statistically different according to the Tukey’s HSD (Honestly Significant Difference) test (p ≤ 0.05).

Ten days after the last treatment, across the three scenarios of infection, treatment with M01 determined markedly low occurrence of disease symptoms, with mean values comparable to those of treatments with the fungicide (F01) (Figure 2 – a2, b2, and c2); however, except for the scenario with infection during treatment (Figure 2 - b2), in which M01 determined a statistical reduction of disease severity comparable just to the fungicide (F01), when pathogen’s infection occurred both before and after treatment, M01 determined an occurrence of disease symptoms statistically similar to that of treatment with RI01 (Figure 2 a2 and c2). After fourteen days, across the three scenarios of infection, treatment with M01 determined an occurrence of disease symptoms (%) significantly lower than untreated control and treatments with P01 and R01 (Figure 2 – a3, b3 and c3); additionally, in the scenario with infection before the treatment (Figure 2 – a3), the application of M01 determined a value of occurrence of disease symptoms as significant as that determined by treatment with F01.

#### 3.1.3. Effectiveness of *Candida oleophila* strain O on lemons

Results of tests on lemons are summarized in Fig.3. As observed previously for clementine tangerine and orange fruit seven days after the last treatment, no lemon fruit from any treatment or infection scenario showed any symptoms of disease (Figure 3 - a1, b1, and c1). Ten days after the last treatment, fruit treated with M01 exhibited a significant low occurrence of disease symptoms, with mean values comparable to those of fruit treated with the fungicide (F01) in scenarios of both infection before and during treatment (Figure 3 – a2 and b2); additionally, treatment with M01 resulted in the lowest occurrence of disease symptoms in the post-treatment infection scenario (Figure 3 – c2). After fourteen days of incubation, when fruit were inoculated before treatment (Figure 3 – a3), the effectiveness of M01 was quite low, with a value of occurrence of disease symptom comparable to that determined by RI01. In contrast, M01 demonstrated strong effectiveness in scenarios with infection both during and after treatment. Specifically, treatment with M01 resulted in the lowest occurrence of disease symptoms in the scenario of infection during treatment (Figure 3 – b3). Additionally, a significant low occurrence of disease symptoms by treatment with M01 was recorded in the scenario with infection after treatment, with a mean value comparable to that of treatment with fungicide (F01) (Figure 3 – c3).

**Figure 3.**
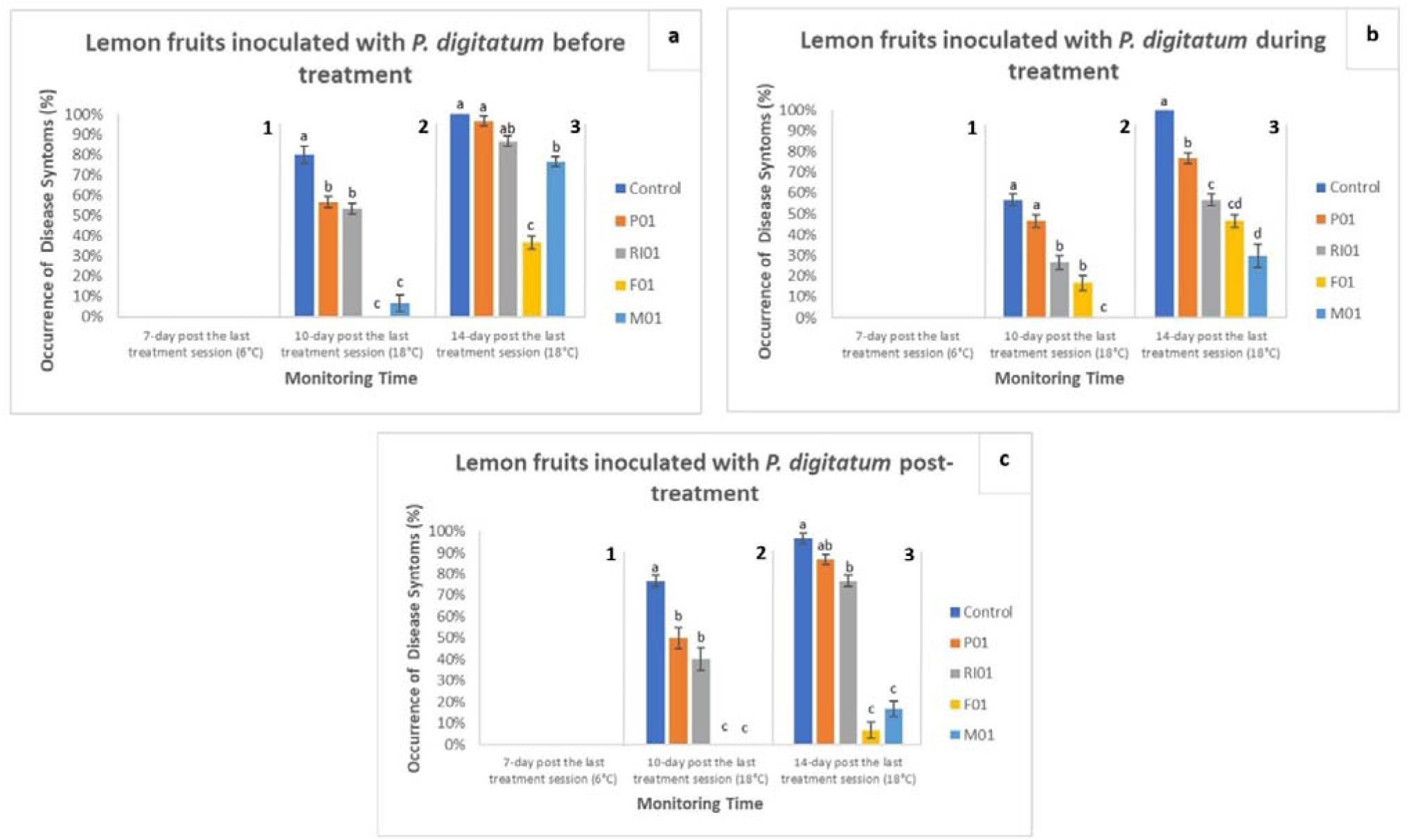
Evaluation of occurrence of disease symptoms (%) in lemon [*Citrus x limon* (L.) Burm. f., cv. Femminello] fruits in three scenarios of infection by *Penicillium digitatum* strain P1PP0: (a) before treatment; (b) during treatment and (c) 12 hours post-treatment. For each treatment, the occurrence of disease symptoms (%) was recorded at 7 (1), 10 (2) and 14 (3) days post the last treatment session. For each time point of measurement, values sharing the same letters are not statistically different according to the Tukey’s HSD (Honestly Significant Difference) test (p ≤ 0.05).

### 3.2. Regulation of induced resistance-related genes in *C. oleophila*-treated citrus fruit

Following paragraph describes the expression levels of the plant defense-related genes, β-1,3-glucanase*-*, phenylalanine ammonia-lyase (PAL)-, and peroxidase (PEROX)-encoding genes measured in peels of clementine tangerine, orange, and lemon fruit at 0-, 24- and 48-hours post after the treatment with M01 and the other three products (Table 1).

Regarding clementine tangerine fruit, upregulation in the levels of β-1,3-glucanase-, PAL-, and PEROX-encoding genes was observed at all time points in peels treated with M01 (Figure 4). Specifically, the treatment with M01 induced the strongest upregulations of the β-1,3-glucanase-encoding gene at 0- and 48-hours post-treatment, while the upregulation recorded at 24 hours was similar to that induced by treatment with F01 (Figure 4a). Concerning the PAL-encoding gene, treatment with M01 showed an increasing trend of upregulation within the first 24 hours; at 48 hours, treatment with M01 still resulted in the strongest upregulation, albeit with a mean value lower than that observed at 24 hours post-treatment (Figure 4b). An increasing trend of expression level was also determined by treatment with M01 towards the PEROX-encoding gene, with the strongest upregulation recorded at 48 hours post-treatment (Figure 4c).

**Figure 4.**
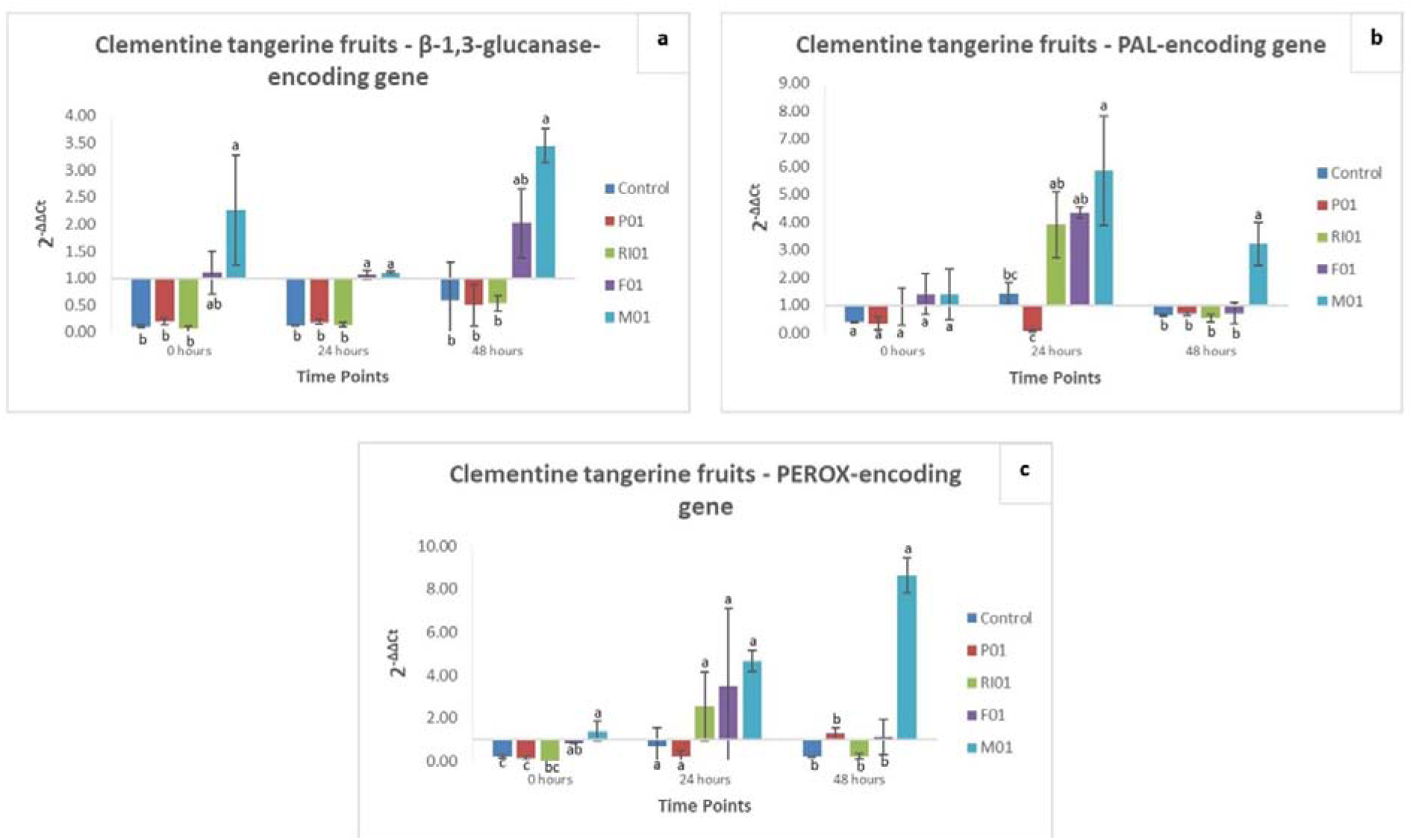
Differences in the expression levels of β-1,3-glucanases-(a), PAL-(b), and PEROX-(c) encoding genes in clementine tangerine fruits (*Citrus clementina* Hort. ex Tan., cv. Comune), wounded and treated with sterile distilled water (controls) or test substances P01 (a sanitization product containing hydrogen peroxide and acetic acid) at 0.1 mL/L, RI01 (a potassium phosphonate formulation) at the dosage of 0.25%, F01 (emulsifiable 50% w/v concentrate formula of imazalil fungicide) at 1.000 ppm (a.i.) and M01 (*Candida oleophila* strain O) as such; at 0, 24, and 48 h after treatment. Values sharing the same letters are not statistically different according to the Tukey’s HSD (Honestly Significant Difference) test (p ≤ 0.05).

With reference to orange fruit, no up-regulation for any gene was induced by any treatment at 0 hours after treatment (Figure 5a, b and c). At 24- and 48-hours post-treatment, an increasing trend of up-regulation was observed for both β-1,3-glucanase- and PAL-encoding genes in fruit treated with F01 and M01, with the highest values in this latter treatment (Figure 5a and b). Finally, the PEROX-encoding gene was up-regulated just in the treatment with M01 and exclusively at 48 hours post treatment (Figure 5c).

**Figure 5.**
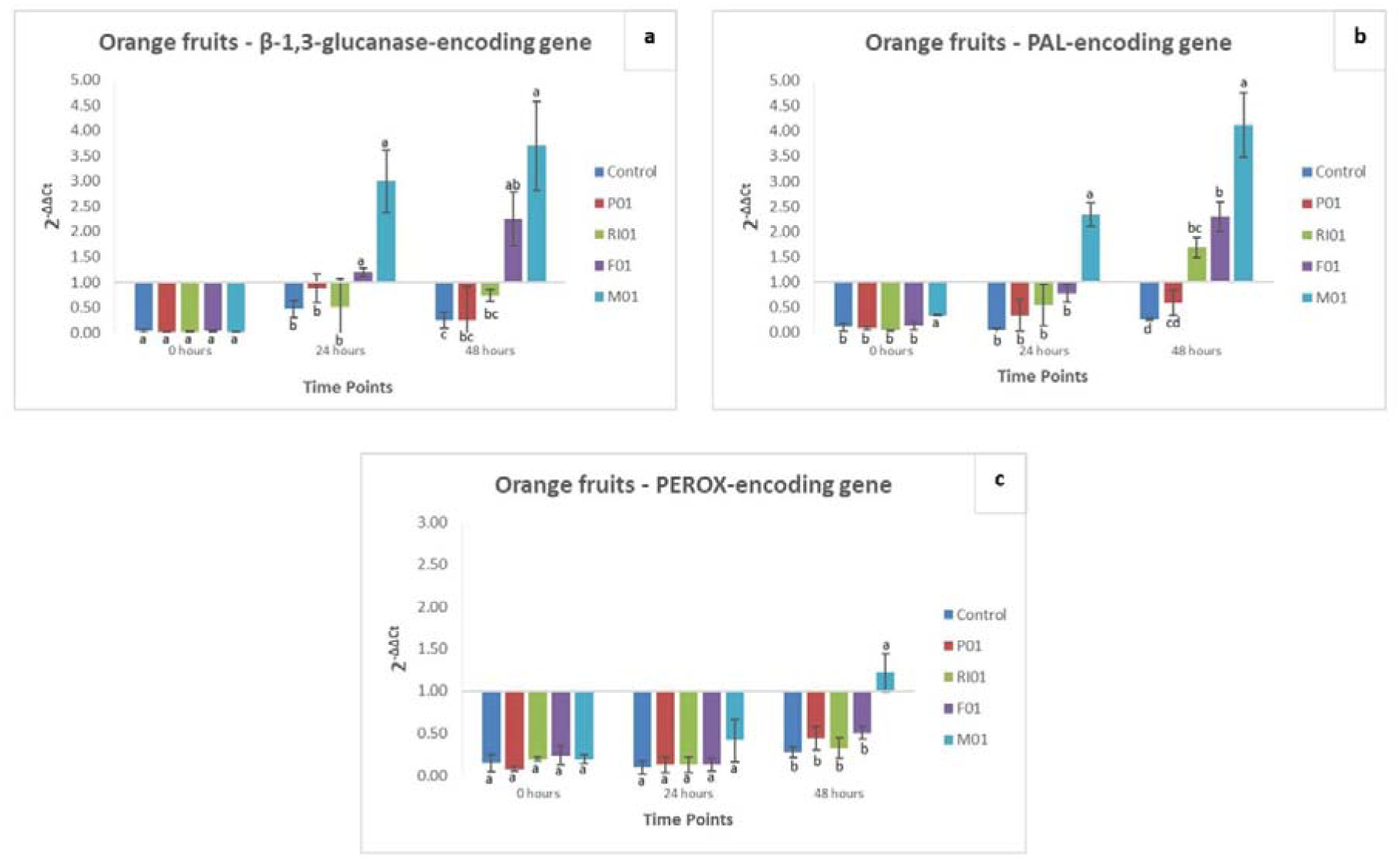
Differences in the expression levels of β-1,3-glucanases-(a), PAL-(b), and PEROX-(c) encoding genes in orange fruit (*Citrus x sinensis*) cv. Ovale, wounded and treated with sterile distilled water (controls) or test substances P01 (a sanitization product containing hydrogen peroxide and acetic acid) at 0.1 mL/L, RI01 (a potassium phosphonate formulation) at the dosage of 0.25%, F01 (emulsifiable 50% w/v concentrate formula of imazalil fungicide) at 1.000 ppm (a.i.) and M01 (*Candida oleophila* strain O) as such; at 0, 24, and 48 h after treatment. Values sharing the same letters are not statistically different according to the Tukey’s HSD (Honestly Significant Difference) test (p ≤ 0.05).

Regarding lemon fruit, no up-regulation for any gene was induced by treatments until 48 hours (Figure 6a, b and c). At this time point, both the β-1,3-glucanase- and PEROX-encoding genes were up-regulated only in the treatment with M01 (Figure 6a and c). Finally, except for the control, up-regulations of PAL-encoding gene were recorded in all treatments, with the highest value in fruit treated with M01 (Figure 6b).

**Figure 6.**
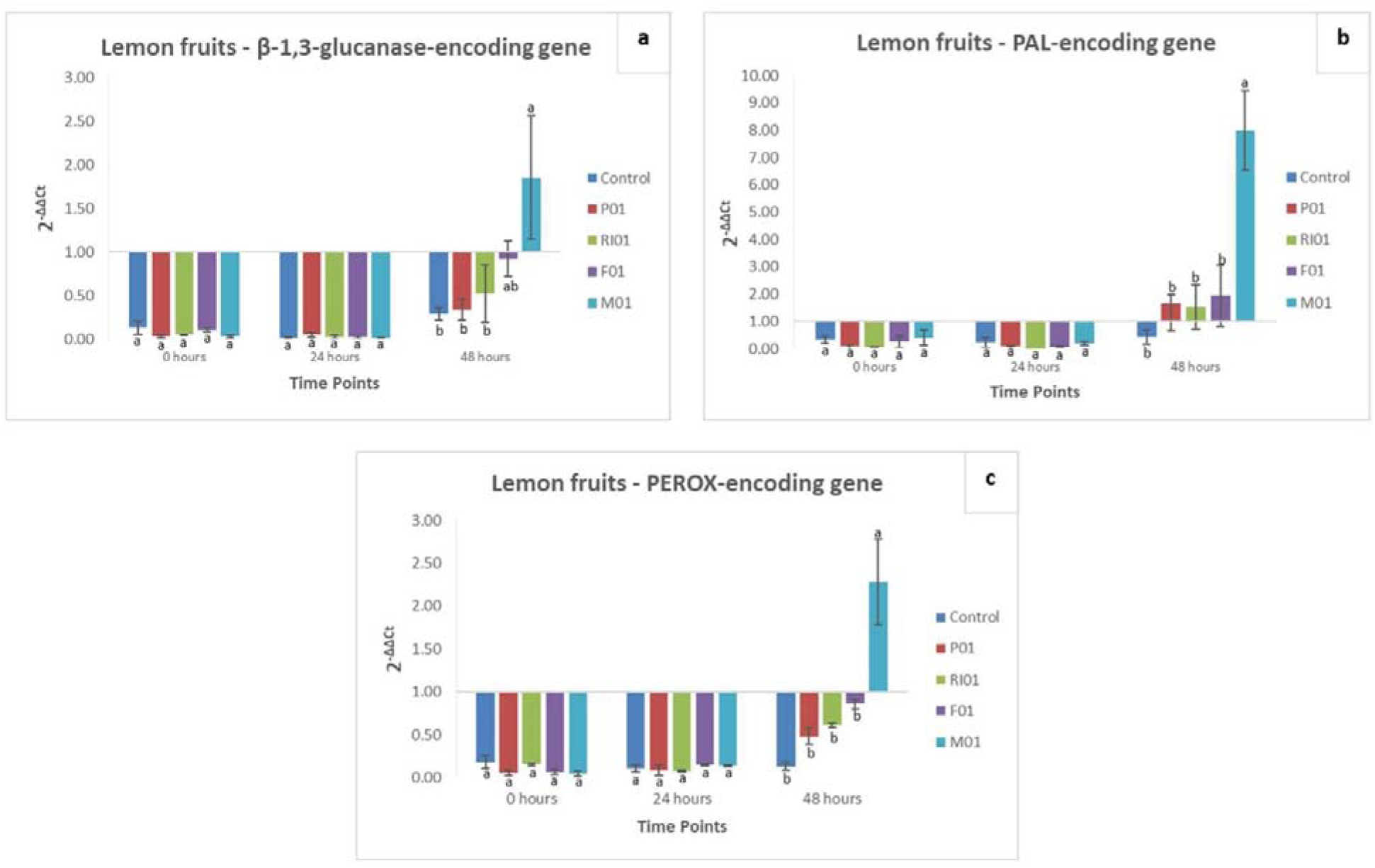
Differences in the expression levels of β-1,3-glucanases-(a), PAL-(b), and PEROX-(c) encoding genes in lemon fruits (*Citrus x limon* L. *Burm. f.,* cv. Femminello), wounded and treated with sterile distilled water (controls) or test substances P01 (a sanitization product containing hydrogen peroxide and acetic acid) at 0.1 mL/L, RI01 (a potassium phosphonate formulation) at the dosage of 0.25%, F01 (emulsifiable 50% w/v concentrate formula of imazalil fungicide) at 1.000 ppm (a.i.) and M01 (*Candida oleophila* strain O) as such; at 0, 24, and 48 h after treatment. Values sharing the same letters are not statistically different according to the Tukey’s HSD (Honestly Significant Difference) test (p ≤ 0.05).

## 4. Discussion

Achieving optimal plant disease control without relying on conventional pesticides is currently one of the most stimulating challenges for the modern Plant Pathology (Bhatta, 2022; Dukare et al., 2022; Lahlali et al., 2022; Nega, 2014; Pal and Gardener, 2006). This ambitious goal holds significant relevance in managing post-harvest decays of fruit from the citrus supply chain. The high incidence of product losses due to molds caused by *Penicillium* species makes it imperative to adopt carefully pondered and evaluated strategies of control. Therefore, with the awareness that means of control safe for human health as effective as pesticides are unlikely to be identified, scientific community has the duty to undertake the study of the efficacy of new plant disease control means through evaluative protocols that realistically simulate the phases of the complex framework of post-harvest of citrus fruit (Zhang et al., 2020). Hence, the first step of this study was to examine different scenarios of *P. digitatum* infections in clementine tangerine, orange, and lemon fruit to evaluate the potential of *C. oleophila* strain O, a specimen well-known for its high and reliable antagonistic activity against post-harvest wound pathogens from various crops (Ballet et al., 2016; Bastiaanse et al., 2010; Dukare et al., 2019; Jijakli and Lepoivre, 1993; Lahlali and Jijakli, 2009; Lahlali et al., 2004, 2011; Lassois et al., 2008), in reducing the occurrence of green mold throughout the main stages of the post-harvest supply chain. Previous studies already investigated some aspects related to the efficacy of *C. oleophila* in controlling diseases of citrus fruit incited by *Penicillium* spp. Brown et al. (2000) evidenced limits of *C. oleophila* strain O in controlling *Penicillum* molds of citrus fruit, demonstrating that the success in controlling fruit decay depends on how quickly and well the yeast colonizes injuries to the fruit surface, including minor injuries involving only oil vesicles. Laahlali et al. (2004) highlighted the good efficacy of *C. oleophila* strain O in reducing mold severity in ’clementine’ and ’valencia-late’ orange fruit inoculated with *P. digitatum* and *P. italicum* after 7 days of incubation at 24°C. A recent study of Hammami et al. (2022) tested the efficiency of *C. oleophila* strain L12 in reducing severity of molds by *Penicillium* spp. in orange fruit subjected to different incubation conditions, including both a long time cold storage (30 days at 4°C) and a particular storage condition consisting of 7 days at room temperature and 30 days at 4 °C. Although these studies have provided valuable information about the potential of *C. oleopihla* in controlling post-harvest decays of citrus fruit, the adopted *in vivo* protocols were based on conditions, such us timing and temperature of incubation, that are far from the reality of the post-harvest supply chain of citrus fruit. Indeed, in the majority of post-harvest chains of citrus fruit, the operational process typically involves bringing fruit from the field into the packinghouses (within four hours from the picking), where blemished or damaged fruit are discarded. Then, marketable fruit undergo a series of steps including washing with detergent and/or fungicides, rinsing with fresh water, waxing (optional), drying, sorting, stamping, sizing, and packing. Finally, fruit are either temporarily held (maximum 12 hours) in a cold store (temperature in the range 4-8°C, depending on the kind of citrus fruit) or directly transported to the market (Malik et al., 2004). The transportation is one of the most critical steps of the post-harvest supply chain of citrus fruit (Cao et al., 2019; Chace et al., 1970; Defraeye et al., 2015). It often exposes fruit to mechanical damages due to improper loading/off-loading procedures, reduced sanitation conditions, and temperatures far from an ideal continuous cold storage environment effective in preserving fruit from the development of *Penicillium* molds (Zacarias et al., 2020). These aspects, together with the rapid progression of these fruit molds and the necessity to minimize post-harvest applications of fungicides, have led citrus producers to realize that the best strategy for reducing product losses is to shorten as much as possible the overall timing of the supply chain. Indeed, nowadays the transportation of fruit typically takes about 2-4 hours for shipments destined for local/regional markets, and approximately 2-7 days for shipments headed to national/continental markets. Once received by the retailer, fruit are displayed on the store shelves at a temperature of shelf-life (16-18°C) and usually sold by no more than seven days from the delivery. Therefore, the duration of a post-harvest supply chain for citrus fruit in a market at a national/continental scale ranges from approximately 10 to 14 days from fruit picking.

In this study, the assessment of tests in different kinds of citrus fruit (clementine tangerines, oranges and lemons) inoculated with *P. digitatum* before, during, or after treatment with *C. oleophila*, provided findings that can be directly related to the efficiency of this biological control agent in typical scenarios of infections of the post-harvest chain of citrus fruit, such as picking, processing in the packinghouse, and loading of fruit for transportation. Additionally, the incubation of inoculated fruit for a period of 7 days at 6°C and for further 7 days at 18°C, simulated the cold storage shipment and the shelf-life phases typical of a market at national/continental scale. Therefore, this study offered the unprecedented opportunity to rationally evaluate the potential of *C. oleophila* for controlling green mold by *P. digitatum* in a real supply chain. In accordance with the physiology of *P. digitatum*, whose cardinal temperatures for growth fall within the range of 6-37°C, no citrus fruit from any treatment or infection scenario exhibited any symptoms of green mold after 7 days of incubation at 6°C. As expected, first symptoms of green mold started to occur at the day 10 day of incubation post-treatment, namely at the third day at shelf-life temperature (18°C); at this time point, the treatment with *C. oleophila* reduced strongly the occurrence of symptoms of green mold in all kinds of tested fruit (clementine tangerines, oranges and lemons) and scenario of infection, with values significantly lowers than untreated controls and often comparable to those recorded in fruit treated with the fungicide IMZ. This marked effectiveness of *C. oleophila* persisted until the 14th day of post-treatment incubation, corresponding to the seventh day at shelf-life temperature (18°C), with values of occurrence of disease symptoms still comparable to those attained with IMZ treatments. The significant reduction in the occurrence of green mold symptoms after treatment with *C. oleophila*, particularly evident at shelf-life temperatures, underscores its efficacy in mitigating post-harvest fungal infections. This sustained efficacy is particularly noteworthy given the challenges posed by prolonged storage and transportation periods within the post-harvest supply chain.

Additionally, the comparability of results to those achieved with conventional fungicide treatments, such as IMZ, highlight the practical potential of *C. oleophila* as an alternative or complementary approach for the management of green molds by *P. digitatum* in the post-harvest chain of citrus fruit at national/continental scale.

This study also provided a significant advancement in understanding the potential of *C. oleophila* to trigger the natural defensive response of various citrus fruit (clementine tangerines, oranges, and lemons). To this purpose, the modulation of the transcriptional levels of genes encoding β-1,3-glucanases, phenylalanine ammonia-lyase, and peroxidase in peels of clementine tangerine, orange and lemon fruit was examined at specific time-points after treatment with *C. oleophila* or comparative substances (potassium phosphonate; IMZ). Several studies supported the involvement of metabolism of β-1,3-glucanases, phenylalanine ammonia-lyase and peroxidase enzymes in the activation of defense response of citrus fruit. The β-1,3-glucanases, an enzyme contributing significantly to the degradation of the fungal cell wall (La Spada et al. 2020, 2021; Youssef et al. 2014), is considered a real molecular marker for postharvest physiological disorders of citrus fruit (Sanchez-Ballesta et al., 2008). The high concentration of phenylalanine ammonia-lyase in citrus peels has been positively associated with the triggering of resistance mechanisms against infections by *P. digitatum*; these mechanisms include the synthesis of antifungal metabolites, such as the phytoalexins scoparone and scopoletin (Ballester et al., 2010; Droby et al., 2002; Kim et al., 1991; La Spada et al., 2021; Youssef et al., 2014). The antioxidant enzyme peroxidase participates in the formation of auxin metabolism, lignin, and suberin, cross-linking cell wall components, synthesizing phytoalexins, and metabolizing reactive oxygen species (ROS) (Almagro et al., 2009). In this study, *C. oleophila* was the treatment that elicited the strongest up-regulations in all kinds of test fruit. Specifically, clementine tangerines resulted the most responsive fruit, with marked up-regulations of the three genes in the treatment with *C. oleophila* at each time-point of measurement. Orange fruit were less responsive than clementine tangerines; for these fruit, treatment with *C. oleophila* stimulated the up-regulation of β-1,3-glucanases- and phenylalanine ammonia-lyase-encoding genes starting from 24 hours post-treatment, with an increasing trend until 48 hours; In contrast, peroxidase up-regulation in orange fruit occurred only at 48 hours post-treatment. Finally, the defensive response of lemon fruit was significantly delayed, with up-regulations of the three genes recorded only after 48 hours. Overall, results from gene expression evidenced that *C. oleophila* was the most efficient treatment in the elicitation of defense response of fruit peels. This high efficiency of *C. oleophila* in activating the studied metabolic pathways is consistent with previous literature. Indeed, Droby et al. (2002) evidenced that the treatment of grapefruit with the strain O of *C. oleophila* promoted the biosynthesis of β-1,3-glucanases and phenylalanine ammonia-lyase enzymes, and that their accumulation in fruit peels showed an increasing trend over time. Another study found that *C. oleophila* applications are also able to promote the gradual accumulation over time of peroxidase enzymes (Zheng et al., 2021).

In conclusion, this study addressed the pressing challenge of achieving optimal plant disease control in citrus fruit without relying on conventional pesticides, emphasizing the significance of managing post-harvest decays in the citrus supply chain. The study focused on evaluating the efficacy of the biological control agent *C. oleophila* strain O in controlling green mold caused by *P. digitatum* throughout various stages of the post-harvest supply chain. Results indicated that *C. oleophila* exhibited significant efficacy in reducing the occurrence of green mold symptoms, particularly under conditions simulating the post-harvest supply chain of citrus fruit. This efficacy was evident even at shelf-life temperatures, highlighting the potential of *C. oleophila* as a practical alternative or complementary approach to conventional fungicides. Moreover, the study contributed valuable insights into the molecular aspects of the defensive response of citrus fruit to *C. oleophila* treatment. The up-regulation of genes encoding β-1,3-glucanases, phenylalanine ammonia-lyase, and peroxidase in fruit peels demonstrated the activation of defense mechanisms. Clementine tangerines displayed the most pronounced response, while oranges and lemons exhibited delayed but significant up-regulation of these genes. Overall, the findings supported the potential of *C. oleophila* as biological control agent not only as pathogen antagonist, but also as elicitor of key defense pathways in citrus fruit. The results highlighted the usefulness of this biological control agent and its comparability with conventional fungicides, emphasizing its importance in the context of a post-harvest citrus industry on a national/continental scale.

## Funding

This study was supported by the project “Smart and innovative packaging, postharvest rot management, and shipping of organic citrus fruit (BiOrangePack)” under Partnership for Research and Innovation in the Mediterranean Area (PRIMA) – H2020 (E69C20000130001), the he “Italie–Tunisie Cooperation Program 2014–2020” project “PROMETEO «Un village transfrontalierpour protéger les cultures arboricoles méditerranéennes enpartageant les connaissances» cod. C-5–2.1–36, CUP 453E25F2100118000 and the European Union (NextGeneration EU), through the MUR-PNRR project SAMOTHRACE (ECS00000022). Ermes Ivan Rovetto was supported by a Ph.D. fellowship funded by University of Catania (cycle XXXVI).

## CRediT authorship contribution statement

**Ermes Ivan ROVETTO**: Writing – original draft, Visualization, Software, Investigation, Formal analysis, Data curation. **Federico LA SPADA**: Writing – review & editing, Writing – original draft, Visualization, Supervision, Methodology, Investigation, Data curation, Conceptualization. **Soumia EL BOUMLASY**: Writing – review & editing, Investigation, Data curation. **Sebastiano CONTI TAGUALI**: Writing – review & editing, Investigation, Data curation. **Mario RIOLO**: Writing – review & editing, Investigation, Data curation. **Antonella PANE**: Writing – review & editing, Validation, Supervision, Resources, Project administration, Funding acquisition. **Santa Olga CACCIOLA**: Writing – review & editing, Validation, Supervision, Resources, Project administration, Funding acquisition, Conceptualization.

## Notes

### Competing Interest Statement

The authors have declared no competing interest.

### Summary of Updates

Title of chapter 2.6; Figure 2; Author affiliation

